# A *MEF2C* transcription factor network regulates proliferation of glomerular endothelial cells in diabetic kidney disease

**DOI:** 10.1101/2024.09.27.615250

**Authors:** Ricardo Melo Ferreira, Debora L. Gisch, Carrie L. Phillips, Ying-Hua Cheng, Maansi Asthana, Blue B. Lake, William S. Bowen, Fang Fang, Mahla Asghari, Angela Sabo, Daria Barwinska, Michael J. Ferkowicz, Robert D. Toto, John R. Sedor, Sylvia E. Rosas, Petter Bjornstad, Jeffrey B Hodgin, Charles E. Alpers, Pinaki Sarder, Jonathan Himmelfarb, Jennifer A. Schaub, Viji Nair, Seth Winfree, Timothy A. Sutton, Katherine J. Kelly, The Kidney Precision Medicine Project, Matthias Kretzler, Sanjay Jain, Tarek M. El-Achkar, Pierre C. Dagher, Michael T. Eadon

## Abstract

The maintenance of a healthy epithelial-endothelial juxtaposition requires cross-talk within glomerular cellular niches. We sought to understand the spatially-anchored regulation and transition of endothelial and mesangial cells from health to injury in DKD. From 74 human kidney samples, an integrated multi-omics approach was leveraged to identify cellular niches, cell-cell communication, cell injury trajectories, and regulatory transcription factor (TF) networks in glomerular capillary endothelial (EC-GC) and mesangial cells. Data were culled from single nucleus RNA and ATAC sequencing and three orthogonal spatial transcriptomic technologies for correlation with histopathological and clinical trial data. We identified a cellular niche in diabetic glomeruli enriched in a proliferative endothelial cell subtype (prEC) and altered vascular smooth muscle cells (VSMCs). Cellular communication within this niche maintained pro-angiogenic signaling with loss of anti-angiogenic factors. We identified a TF network of MEF2C, MEF2A, and TRPS1 which regulated SEMA6A and PLXNA2, a receptor-ligand pair opposing angiogenesis. In silico knockout of the TF network accelerated the transition from healthy EC-GCs toward a degenerative (injury) endothelial phenotype, with concomitant disruption of EC-GC and prEC expression patterns. Glomeruli enriched in the prEC niche had histologic evidence of neovascularization. MEF2C activity was increased in diabetic glomeruli with nodular mesangial sclerosis. The gene regulatory network (GRN) of MEF2C was dysregulated in EC-GCs of patients with DKD, but sodium glucose transporter-2 inhibitor (SGLT2i) treatment reversed the MEF2C GRN effects of DKD. The MEF2C, MEF2A, and TRPS1 TF network carefully balances the fate of the EC-GC in DKD. When the TF network is “on” or over-expressed in DKD, EC-GCs may progress to a prEC state, while TF suppression leads to cell death. SGLT2i therapy may restore the balance of MEF2C activity.

## Introduction

Diabetic Kidney Disease (DKD) is the leading cause of Chronic Kidney Disease (CKD), affecting over 12 million individuals in the United States^1^. Early manifestations of DKD include thickening of the glomerular basement membranes and mesangial matrix expansion. As Olson, Heptinstall, and others have noted, podocyte effacement and neovascularization with increased endothelial density materialize as DKD advances^2^. Late stage DKD is characterized by Kimmelstiel-Wilson (KW) nodule formation, podocyte loss, and glomerulosclerosis^3^. While podocyte injury in DKD is well described ^4^, a knowledge gap persists regarding the relationship between nodular mesangial sclerosis and neovascularization.

Manifestations of DKD such as glomerular hypertension, hypoxia, and hyperglycemia all contribute to a complex milieu wherein the glomerular cell composition and communication change in response to stress^5^. Recently, a single-cell RNA sequencing (scRNAseq) study yielded an atlas of renal endothelial cells with key insights into blood pressure regulation^6^. However, glomerular response to stress is heterogeneous^7^, with localized histologic changes. Therefore, the spatial context will augment our understanding of DKD pathophysiology^8^.

Glomerular hypertension is a hallmark of DKD and may result in localized hypoxia^9^. Glomerular pressure is regulated distinctly from systemic blood pressure. Both hemodynamic changes and hormonal signaling drive glomerular hypertrophy and angiogenesis to compensate for hypoxia^9^. Vascular Endothelial Growth Factor A (VEGFA) expression within the glomerulus is necessary to maintain a healthy microvasculature^10^. VEGFA and B are upregulated in early DKD and their loss contributes to endothelial and mesangial cell defects^11^, ultimately exacerbating proteinuria and glomerulosclerosis in murine models^10^; however, inhibition of VEGFB failed to reduce proteinuria in a Phase II trial^12^. A diversity of receptor-ligand pairs has been invoked to regulate cross-talk between the podocyte, endothelial cell, and mesangial cell. These signaling pathways include those initiated by angiopoietin, platelet-derived growth factor-B Chain (*PDGFB*), and transforming growth factor-β^5^.

Recently, we described a case of neovascularization in a patient with DKD^13^. In this case, three-dimensional CD31 protein staining confirmed the multiple afferent arterioles and pericapsular neovessels observed histologically. An increase in endothelial cell density within the glomeruli was accompanied by upregulation of angiogenic gene expression assessed by spatial transcriptomics (ST). We expand upon this study by now interrogating all reference and DKD samples within the Kidney Precision Medicine Project (KPMP) and Human Biomolecular Atlas Program (HuBMAP) sc/snRNA-seq and Visium ST atlas of the kidney^14^. Using deconvolution tactics, we have previously defined colocalizing cell types with their attendant injury signatures and cellular cross-talk^15, 16^. While progress has been made in cataloging proximal tubule cell transitions^17^, including transcription factor (TF) networks that regulate phenotypic alterations in cellular response to injury^17^, these approaches may also yield insights into the glomerular endothelial cell changes of DKD to define patient subgroups for targeted therapy. In this study, we hypothesized that spatial localization and cell communication would underlie the endothelial and vascular smooth muscle cell signature alterations associated diabetic kidney disease. To this end, we utilized orthogonal snRNA-seq and multiome atlases with three spatial transcriptomic datasets to define a proliferative endothelial cell with its associated changes in glomerular cell composition, expression trajectories, localization, communication, and regulation in the histologic context of DKD.

## Methods

### Sample acquisition

For Visium fresh frozen spatial transcriptomics, kidney biopsy samples were obtained with informed consent through the KPMP consortium (https://KPMP.org) and approved under a single Institutional Review Board (IRB #20190213) protocol. Formalin-fixed paraffin-embedded (FFPE) samples were obtained under waiver of informed consent from the Biopsy Biobank Cohort of Indiana (BBCI)^18^, approved by the Indiana University IRB (#1906572234), and by the Kidney Translational Research Center (KTRC) as part of the HuBMAP consortium under IRB #201102312 at Washington University in St. Louis. Samples were classified as DKD according to the clinical diagnosis at the time of biopsy (Supplemental Table S1).

### Experiment and analysis of omics technologies

Kidney tissue samples underwent Visium Spatial Transcriptomics (ST)^19^; Visium FFPE Spatial Transcriptomics (FFPE); and Xenium *in situ* sequencing (ISS) with a custom probe set (BMU84Y, Supplemental Table S2). Complementary disaggregated cell technologies include single nucleus RNA sequencing (snRNAseq), obtained from the published KPMP-HuBMAP atlas^14^, and multiome data obtained from the KPMP-HuBMAP epigenetic atlas^17^. For downstream analysis, the following algorithms were utilized. Nuclei were clustered correcting for batch effect with Harmony 1.2.0^20^. Cell-cell communications were predicted in the snRNAseq glomerular nuclei using CellChat 2.1.2^21^. Visium raw reads were normalized with SCTransform for batch effect correction^16^. To estimate cell type composition in Visium Spatial Transcriptomics (ST), the expression of each spot is deconvolved using transfer scores from atlas annotations, calculated using the anchors’ integration method from Seurat 4.3.3^22^. To identify cells in ISS, a subset of the snRNAseq dataset was mapped with the anchors’ method of Seurat 5.0.3. Pathway enrichment for snRNAseq clusters was performed with SCPA 1.6.1^23^. Cellular niches were defined by clustering ST spots according to their cell type composition using traditional Louvain clustering and UMAP projection methods from Seurat. A Fisher’s exact test was utilized to evaluate the likelihood of each cellular niche being overrepresented in DKD samples. We applied the Seurat anchors method in the reverse direction to back-map the ST niche annotations to the glomerular cells of the multiome dataset to identify a proliferative endothelial (prEC) subgroup preferentially mapping to the DKD endothelial ST niche. In the multiome dataset, peaks were annotated by MACs2, and JASPAR2022 (v.0.99.7) was used as the motif database. We employed scMEGA (v.1.0.2) to model cell trajectories and regulatory interactions^24^. The *in silico* knockout was modeled with CellOracle (v.0.12.0)^25^. Individual glomeruli in the Visium FFPE dataset were annotated for histopathological features relevant to DKD, and association with neighborhoods or MEF2C activity accessed with a Fisher’s test. Expression of genes regulated by MEF2C was evaluated in an independent human kidney scRNAseq dataset^26^ which included DKD patients treated with SGLT2i. For further details on the experiments and analysis, see Supplemental Methods.

## Results

### Subjects

Nodular mesangial sclerosis and neovascularization are common manifestations of DKD. To elucidate the cellular cross-talk underlying the spectrum of DKD within the glomerulus, we interrogated orthogonal data derived from a partially overlapping set of kidney samples (Supplemental Table S1). The total number of human kidney specimens interrogated was 74 including snRNAseq (N= 31), the multiome combined snRNA-seq and scATAC-seq (N= 12), 10x Visium fresh frozen spatial transcriptomics (N= 37 samples [14 reference, 23 DKD], 31545 spots, 512 glomeruli), FFPE ST (N= 12 samples [8 reference, 4 DKD], 22052 spots, 264 glomeruli), and Xenium *in situ* sequencing (ISS, N= 7 samples [4 reference, 3 DKD], 40551 glomerular cells, 236 glomeruli).

### Spatial mapping of glomerular cell types in diabetic kidney disease

To examine the glomerular milieu in DKD, we extracted podocytes, endothelial, and mesangial cells / VSMCs from the KPMP-HuBMAP snRNA-seq kidney atlas^14^ (Figure 1A). Comparing healthy reference and DKD samples, expected pathways^23^ were enriched in glomerular cell types (Figure 1B). For podocytes, enriched pathways included development and adhesion, sphingolipid metabolism, and VEGF signaling - all associated with mesangial expansion and neovascularization in DKD^27^. Endothelial cell pathways were enriched for proliferation and migration, consistent with the observed angiogenesis in DKD^28–30^. VSMC pathways aligned with the processes described above, including development, angiogenesis, and signaling. To quantitate cell type changes in DKD, we histologically annotated glomeruli of ST samples to differentiate glomerular and tubulointerstitial spots. Then, we deconvoluted the cell type composition of 1877 glomerular spots using a snRNAseq label transfer methodology^31^ (Figure 1C). Across all glomeruli, the proportion of podocyte signature decreased in DKD with an increase in endothelial and VSMC signature relative to reference (Figure 1D). These changes are consistent with podocyte effacement, neovascularization, and mesangial expansion often observed in DKD^9^. A similar cellular composition was observed with ISS, confirming upregulation of endothelial and mesangial cells within the diabetic glomeruli (Figure 1E).

**Figure 1.**
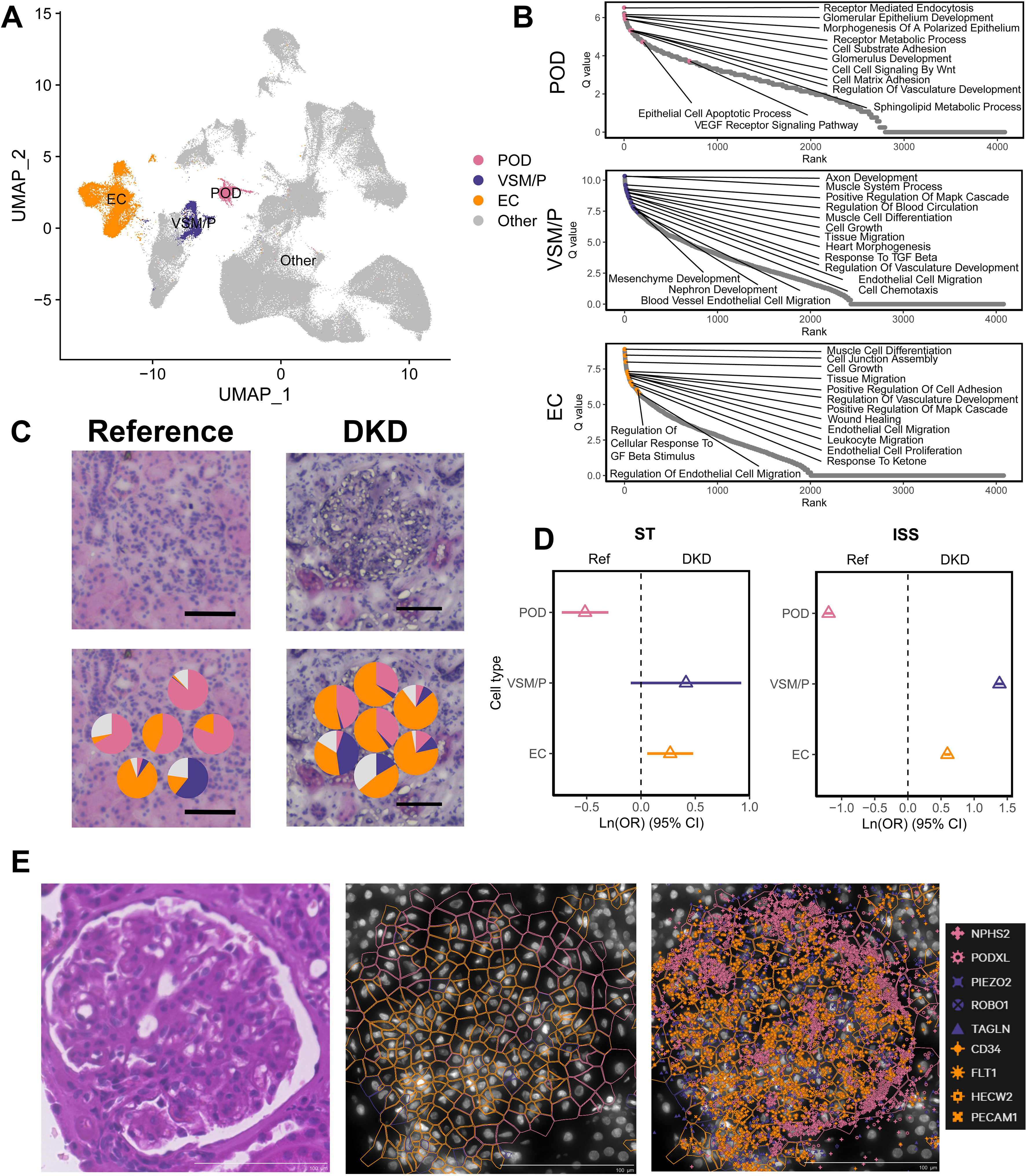
Glomerular cell types in diabetic kidney disease: **A)** A glomerular snRNAseq object was constructed with 16,488 nuclei representing cell types potentially localizing to glomeruli including all podocyte (POD), all endothelial cell (EC), and all vascular smooth muscle and mesangial cell (VSM/P) subtypes. **B)** Pathways enriched in the podocyte, VSM/P, and EC were determined for diabetic kidney disease (DKD, N=7) and reference (Ref) samples (N=18). **C)** Examples of Visium glomerular mapping of cell types in reference and DKD. An increased proportion of endothelial cell signature (orange) is observed in the diabetic glomerulus. Scale bar = 100 µm. **D)** The odds ratio of cell type signature vectors localized to glomeruli of reference and DKD samples in Visium ST (N=36). DKD glomeruli had reduced podocyte signature and increased EC signature as measured by Visium ST. For Xenium in situ sequencing (ISS, N=7), the number of cells of each cell type were localized to glomeruli of reference and DKD samples. VSM/P and EC cells were more abundant in DKD glomeruli, while podocytes were less frequent. **E)** An example of cell type mapping and segmentation with ISS, with expected markers defining each cell type. Pink – podocyte, Purple – VSM/P or mesangial cell, Orange – endothelial cell. Scale bar = 100 µm.

### Niche analysis uncovers a proliferative endothelial cell subtype

To define the spatial distribution of healthy and injured cell types as niches, we selected KPMP atlas-defined glomerular cell types including podocytes (POD), glomerular capillary endothelial cells (EC-GC), injured / degenerative Endothelial Cells (dEC), mesangial Cells (MC), injured / degenerative VSMCs (dVSMCs), other endothelial cells (now annotated as “not specified” or EC-NS) and other VSMCs (VSMC-NS) (Figure 2A). Expected marker gene expression was observed for POD, EC-GC, and MC, while injury genes were expressed in degenerative cell types (Figure 2B). Both EC-NS and VSMC-NS had reduced gene expression of known markers of EC-GCs (*e.g. HECW2*) and MCs (*e.g. ROBO1*). Upon mapping these cell types onto glomerular spots, an increase in endothelial cell signature was observed in DKD, driven by increased EC-NS, rather than EC-GC or dEC signature (Figure 2C). Although the EC-NS clusters were not originally annotated as glomerular cell types in the KPMP-HuBMAP atlas, EC-NS localized to glomerular spots proportionately more than TI spots. The glomerular mapping of the EC-NS and its proportional increase in DKD provided an impetus to identify an endothelial cell subtype, potentially associated with neovascularization.

**Figure 2.**
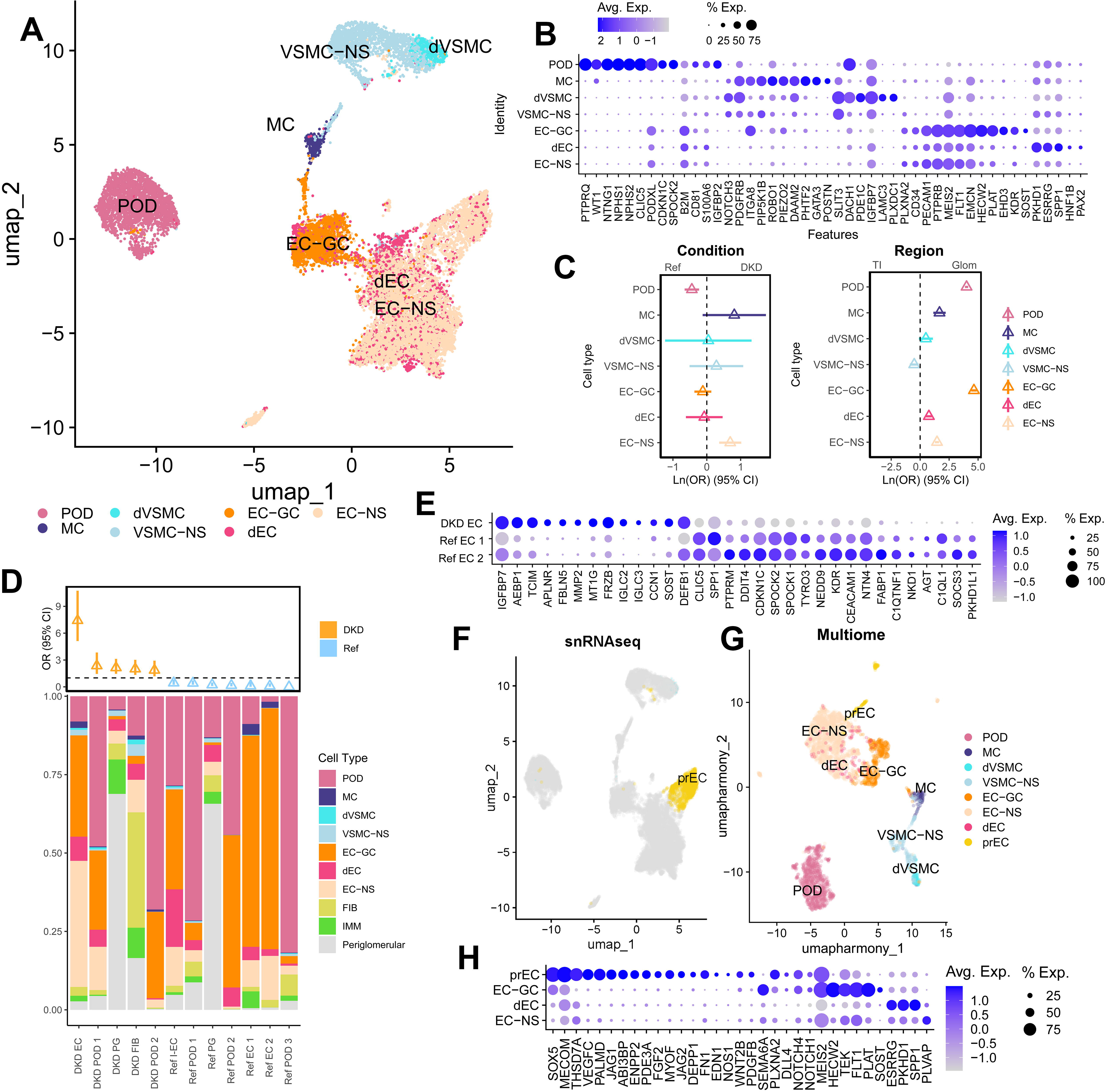
Niche localization of proliferative endothelial cell subtype: **A-B)** Podocytes (POD), endothelial cells (EC) and vascular muscle cells (VSMC) were reclustered and annotated. Expression patterns within glomerular capillary endothelial cells (EC-GCs) and degenerative / injured endothelial cells (dEC) were compared to a composite cluster of all other endothelial cells (not specified or EC-NS). Mesangial cells (MCs), degenerative or injured VSMCs (dVSMC), and the remaining pool of vascular smooth muscle cells (VSMC-NS) were similarly compared. **C)** The glomerular mapping proportion from Visium ST according to condition (reference/Ref or diabetic kidney disease/DKD) and region (tubulointerstitium [TI] or glomerulus [glom]) revealed an increase in EC-NS signature in DKD glomeruli, suggesting an important unidentified glomerular cell type was embedded within the EC-NS cluster. Other cell types mapped to the appropriate region as expected (e.g. POD and MC in the glomerulus). **D)** All Visium glomerular ST spots (N= 1877) were clustered according to cell type vector proportion to define niches of colocalizing cells. Of the 21 identified niches, twelve were significantly enriched in either Ref or DKD samples, including a DKD EC niche with enrichment of EC-NS signature. **E)** Expression of selected genes differentially expressed between endothelial niches localizing to DKD or reference. **F-G)** The DKD EC niche signature was back-mapped into the glomerular snRNAseq and multiome object to identify a unique subcluster of endothelial cells defined by a proliferative or pro-angiogenic signature (prEC). **H)** Expression of selected genes differentially expressed in prEC in relation to other endothelial cells (prEC markers) in snRNAseq.

To characterize colocalization niches, we clustered glomerular spots according to cell type composition (Supplemental Figure S1), identifying niches enriched in DKD or reference samples (Figure 2D). For example, a fibroblast-dominant niche (DKD Fib) was enriched in DKD. Three endothelial-dominant niches were identified, one in DKD (DKD EC) and two in reference (Ref EC 1, Ref EC 2). In the Ref EC 1 and 2 niches, EC-GC signature predominated, representing niches with healthy glomerular capillaries. In contrast, the DKD EC niche consisted of a high proportion of EC-NS. A differential expression analysis comparing DKD EC spots to Ref EC 1 and 2 spots identified genes associated with angiogenesis (*IGFBP7, CCN1, MT1G*), signaling (*TCIM, FRZB, SOST*), and injury (*MMP2, DEFB1*) (Figure 2E). We then back-mapped the expression signature of the three niches onto the snRNAseq and multiome atlases (Supplemental Figure S2) to identify the subcluster of EC-NS relevant to the DKD EC niche. The expression signature of this EC-NS subcluster is consistent with a unique DKD proliferative endothelial cell (prEC) phenotype (Figure 2F-G). The prEC could be further subtyped based on afferent arteriole (*PTGS1*), EC-GC (*MMP2*), and vasa recta (*ENPP2*) injury markers suggesting an anatomical trajectory embedded within this subtype (Supp. Figure 2). Thus, prECs is a subcluster of EC-NS that localizes within DKD glomeruli. Differential expression between prECs and other endothelial cells in snRNAseq identified marker genes associated with angiogenesis (*SOX5, VEGFC, PALMD, ENPP2, MECOM, PDGFRB, JAG1)*, with the absence of known EC-GC (*HECW2, TEK*) and injury (*ESRRG, SPP1*) markers (Figure 2H).

### Anti-proliferative communication through SEMA6A and PLXNA2 is reduced in DKD

We sought to understand the cellular cross-talk underlying the changes in cell type proportion within DKD and reference niches. Using snRNAseq, we modeled communication between cell types with known receptor-ligand (RL) pairs in reference and DKD glomeruli (Figure 3, Supplemental Table S3). RL pairs associated with angiogenesis and cell proliferation were enriched in EC subtypes (Figure 3A). The *VEGFA*–*VEGFR* axis, a ligand-receptor pair vital to maintaining healthy glomerular vasculature^27^, was present in both reference and DKD glomeruli. In contrast, the *SEMA6A*-*PLXNA2* RL pair, postulated to inhibit angiogenesis, was lost in certain endothelial cells of DKD. *SEMA6A* (Semaphorin 6a), expressed by endothelial cells, and *PLXNA2* (Plexin A2), expressed by both endothelial and VSMCs, are known to regulate cell adhesion^32^, morphogenesis^33, 34^, and to modulate *VEGFA* signaling with an antiangiogenic effect^35,36^. Using ISS, we determined that glomerular expression density was reduced for *SEMA6A* (relative to *PECAM1*) and *PLXNA2* (relative to the sum of *PECAM1* and *TAGLN*) in DKD (Figure 3B).

**Figure 3.**
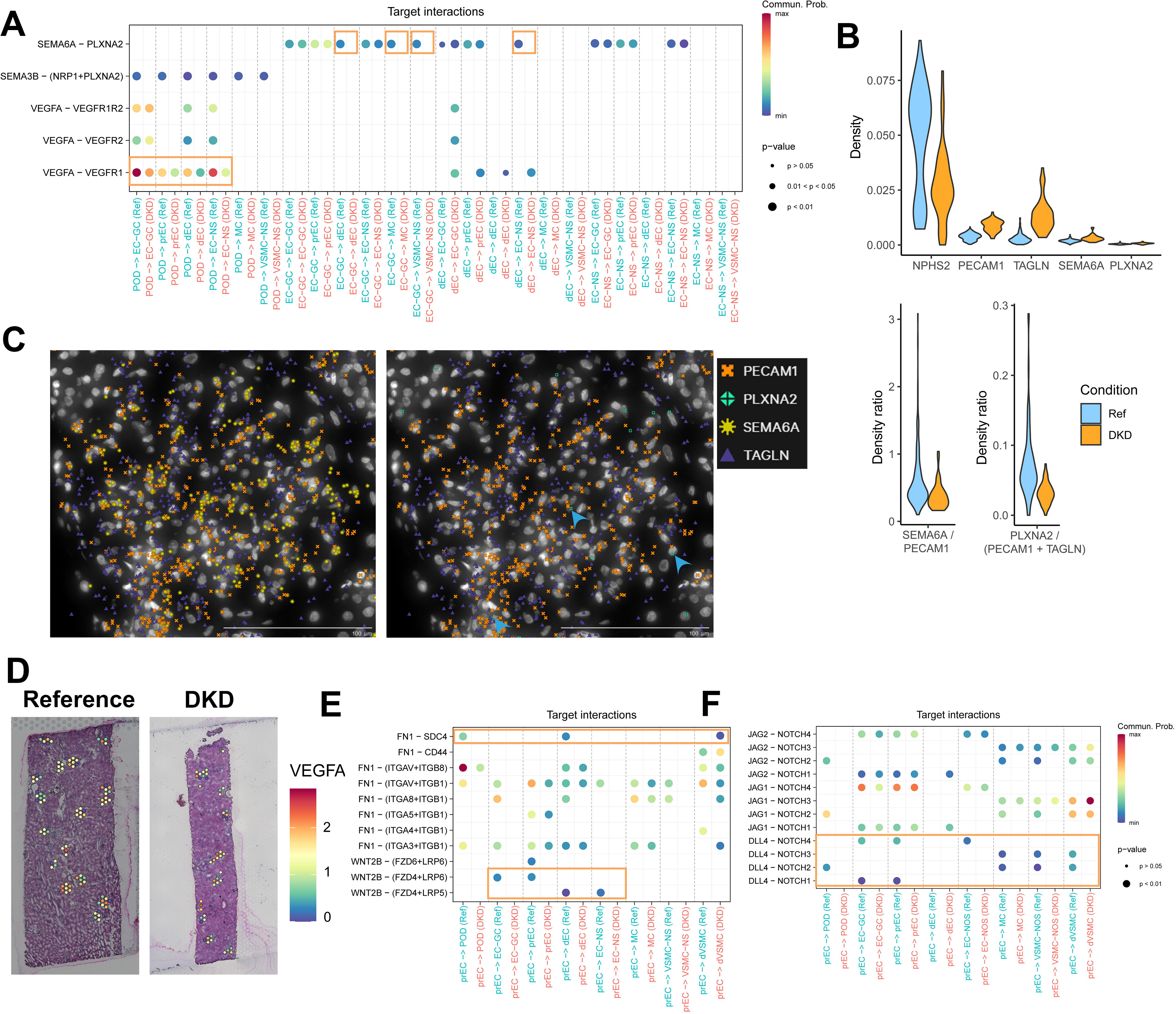
Glomerular cell-cell communication: **A)** Angiogenesis-associated communication stemming from podocytes (POD) or endothelial cells (ECs). **B)** Transcript density of POD, EC, and mesangial cell (MC) markers from glomeruli of in situ sequencing samples. The density ratios of *SEMA6A* to *PECAM1* and *PLXNA2* to *PECAM1+TAGLN* were reduced in diabetic kidney disease (DKD). **C)** The colocalization of *SEMA1* (left) and *PLXNA2* (right) in a glomerulus. Blue arrowheads point to nuclei with colocalization of *PLXNA2* and either *PECAM1* or *TAGLN*. **D)** *VEGFA* expression localizing over glomeruli in a reference and DKD sample. **E)** Angiogenesis-associated communication stemming from proliferative endothelial cells (prECs). **F)** Jag-Notch and *DLL4*-Notch communication involving proliferative endothelial cells. Glomerular capillary endothelial cells: EC-GC; degenerative endothelial cell: dEC; proliferative Endothelial cell: prEC; endothelial not specified (without prEC): EC-NS; mesangial cells: MC, degenerative vascular smooth muscle cell (dVSMC); VSMC not specified: VSMC-NS.

To spatially contextualize these findings, we observed that *SEMA6A* and *PLXNA2* appropriately colocalize with *PECAM1* and *TAGLN*, in endothelial and VSMCs respectively (Figure 3C). *VEGFA* expression was maintained in reference and DKD samples (Figure 3D).

Other proangiogenic RL signaling was observed in DKD. Specifically, dECs expressed *VEGFA* in DKD, potentially contributing to EC proliferation. The prEC exhibited a pro-proliferative communication pattern with other endothelial cells in DKD through the RL pair *WNT2B*-*FZB4*^37, 38^ (Figure 3E). We found dysregulation of RL pairs involving fibronectin (*FN1*), involved in wound healing and vascular development^39, 40^. Finally, prECs also exhibited a unique communication pattern through the Notch (Figure 3F) pathway. The *DLL4*-Notch pair inhibits angiogenesis^41^ and regulates proliferation and branching of new vessels^42^. prECs have reduced expression of the *DLL4* receptor in DKD, while *JAG1*/*JAG2*-Notch communication was maintained. Similar to the *VEGFA*-*VEGFRA* RL pair, angiopoetin-1 (*ANGPT1*) and TIE-2 (*TEK*) communication was maintained across all EC subtypes and in the MC and dVSMC. In contrast, other known RL pairs were dysregulated, including upregulation of platelet-derived growth factor-B Chain (*PDGFB*) in prECs and its receptor (*PDGFRB*) in podocytes and all MC / VSMC subtypes. Likewise, transforming growth factor-β (*TGFB1*) signaling through its receptors (*TGFBR1, TGFBR2*) was upregulated in prECs and dVSMCs (Supplemental Table S3).

### MEF2A, MEF2C, and TRPS1 control endothelial cell state transitions

We then leveraged a multiome dataset to define the regulatory networks underlying endothelial cell state transitions. Upon subsetting EC-GC and dEC cells (Figure 4A), we identified a trajectory emanating from the EC-GC (Figure 4B) and modeled the gene regulatory network (GRN) controlling this trajectory (Supplemental Table S4). The TFs *MEF2A, MEF2C,* and *TRPS1* present high centrality (Figure 4C) and are predicted to regulate *SEMA6A* and *PLXNA2* (Figure 4D). Within the multiome, we identified open chromatin peaks where these TFs are predicted to bind (Figures 4E-G) with reduced accessibility in dEC as compared to EC-GC. The mRNA expression is consistent with the accessibility. *MEF2A* and *MEF2C* are known to be involved in the maintenance of endothelial cell function and regulation of angiogenesis^43–47^. TRPS1 is implicated in angiogenic manifestations in tumors^48, 49^. *MEF2A, MEF2C,* and *TRPS1* also regulated a trajectory for the EG-GC to prEC transition (Figure 4H-J), with the prEC marker gene, *PDGFB*, positively regulated by this TF network (Supplemental Figure S3)

**Figure 4.**
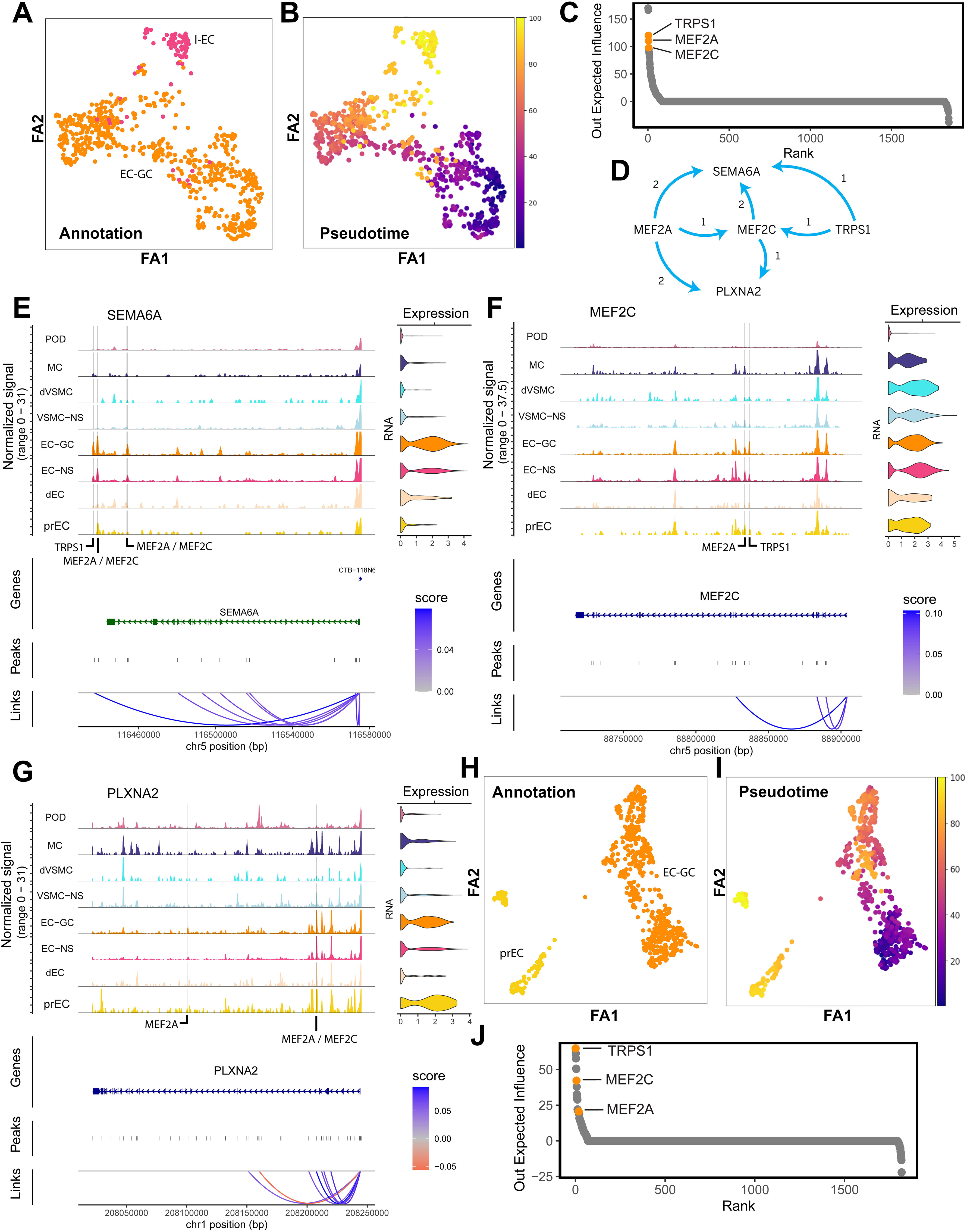
Transcription factor regulation of endothelial cell trajectories: **A)** Force directed projection of Endothelial Cell Glomerular Capillaries and degenerative endothelial cells (dEC). **B)** Pseudotime trajectory from EC-GC to dEC. **C)** Out influence centrality measure of the gene regulatory network associated with the EC-GC to dEC trajectory identifies *MEF2A*, *MEF2C*, and *TRPS1* as a transcription factor network. **D)** *MEF2A*, *MEF2C*, and *TRPS1* regulate *SEMA6A* and *PLXNA2* (number of peaks regulated is stated next to arrows). **E)** Chromatin accessibility tracks of *SEMA6A* for glomerular cell types with predicted accessible binding sites of *MEF2A*, *MEF2C*, and *TRPS1*. **F)** Chromatin accessibility tracks of *MEF2C* for glomerular cell types with predicted accessible binding sites of *MEF2C* and *TRPS1*. **G)** Chromatin accessibility tracks of *PLXNA2* for glomerular cell types with predicted accessible binding sites of *MEF2A* and *MEF2C*. **H)** Force directed projection of Endothelial Cell Glomerular Capillaries and proliferative endothelial cells (prEC). **I)** Pseudotime trajectory from EC-GC to prEC. **J)** Out influence centrality measure suggests the same gene regulatory network controls the EC-GC to prEC trajectory.

### Transcription factor knock-out disrupts endothelial cell trajectories

To evaluate the effect of *MEF2A, MEF2C,* and *TRPS1* in the glomerular endothelium, we performed *in silico* combined knock-out (KO) of the three TFs for two trajectories: EC-GC to either the dEC or prEC. CellOracle estimated the previously identified GRN in subclusters along the EC-GC to dEC trajectory (Figure 5A). The development flow of individual cells was qualitatively evaluated before and after KO (Figure 5B). The magnitude of developmental changes accelerated as assessed by the internal product of the two vectors in the force directed projection (Figure 5C) and along the pseudotime (Figure 5D). The KO resulted in a decrease of *SEMA6A, PLXNA2,* and *FZB4* in ECs. The individual KO of each TF was qualitatively similar (Supplemental Figure S4). For the EC-GC to prEC trajectory, the combined TF KO is shown to disrupt the trajectory, not accelerate it (Supplemental Figure S5). In summary, the *MEF2A, MEF2C,* and *TRPS1* TF network carefully balance the fate of the EC-GC. When the TF network is “on” or expressed, an EC-GC may progress to a prEC state. When the TF network is off, the EC-GC progresses toward an dEC or cell death state.

**Figure 5.**
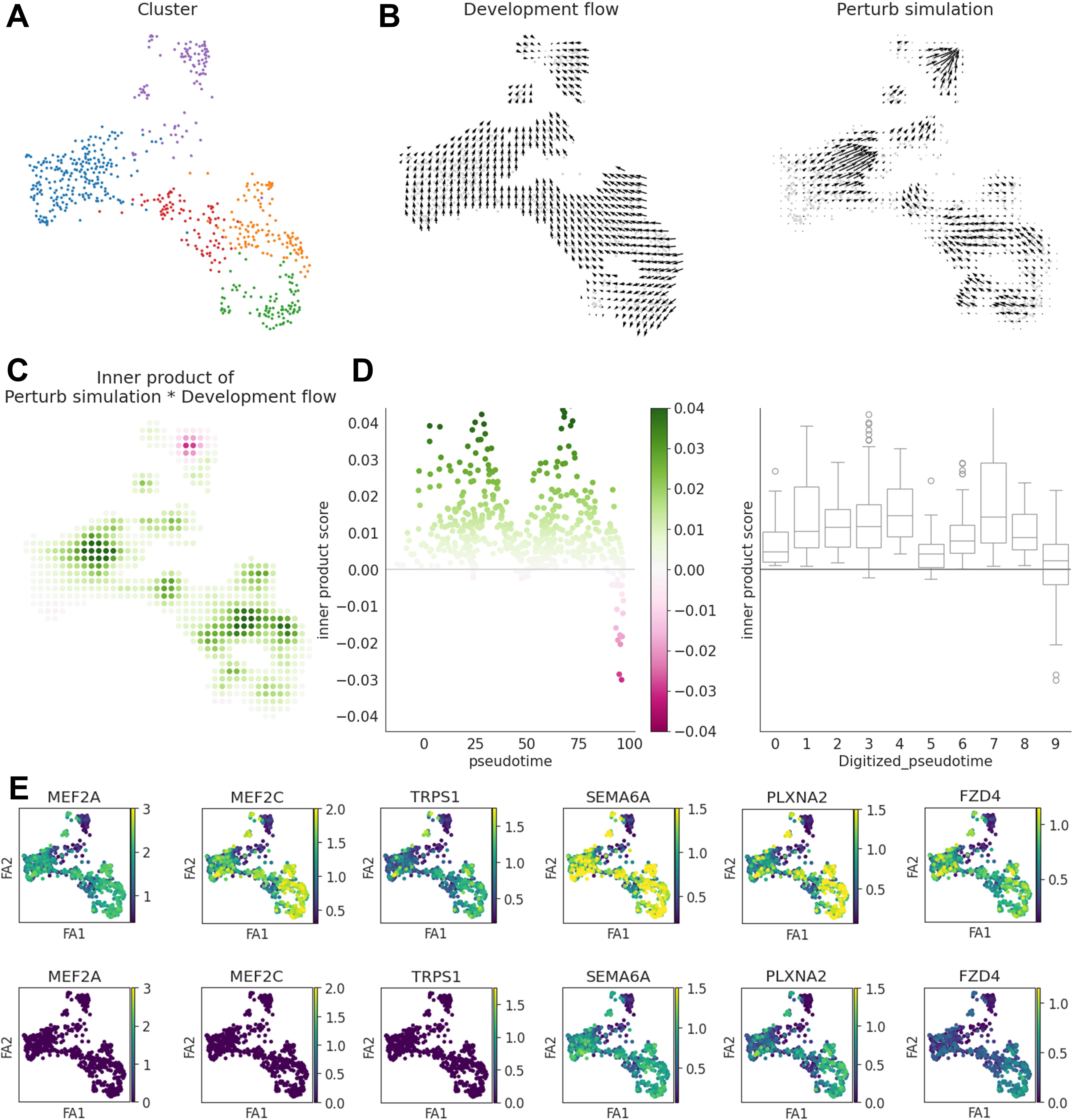
Combined *in silico* knockout: **A)** CellOracle modeled subclusters in the EC-GC to dEC trajectory. **B)** Development flow (left) is accelerated after knockout (right). **C)** Internal product between developmental fields before and after knockout shows acceleration throughout the EC-GC transition to dEC after knockout. **D)** Internal product between developmental fields along the pseudotime. **E)** Expression of transcription factor and target genes before and after knockout reveal reduction of *SEMA6A* and *PLXNA2* after knockout.

### Vascular smooth muscle cell state transitions

A trajectory was identified from MC to VSMC-NS, ending in dVSMC (Supplemental Figure S6 A-B). In the GRN modelled for this trajectory, a TF of the myocyte enhancer factor 2 family, *MEF2D*, was identified to regulate the expression of its target gene *SMAD3* (Supplemental Figure S6 C). The chromatin accessibility in the predicted binding sites is consistent with the mRNA expression. *In silico* KO of this TF disrupted the trajectory directing the flow from MC and dVSMC to the VSMC-NS. (Supplemental Figure S6 D-G).

### Histopathological associations with spatial niches and MEF2C activity

To correlate our molecular findings with histology, we performed ST in FFPE tissue. A trained pathologist scored the 7 µm thick H&E and 3 µm thick PAS sequential sections for each individual glomerulus for these features: hilar neovascularization, non-nodular mesangial sclerosis (mesangial expansion), nodular mesangial sclerosis (Kimmelstiel-Wilson nodule), and segmental glomerular sclerosis (SGS). We then merged the FFPE ST dataset and mapped onto it the niches previously identified. We observed that the endothelial DKD niche (DKD EC) had higher transfer scores than the Ref EC 2 in glomeruli with neovascularization (Figure 6A). The Ref EC 1 niche presented no mapping in FFPE tissue. The pro-fibrotic DKD FIB niche mapped to glomeruli with SGS lesions (Figure 6B).

**Figure 6.**
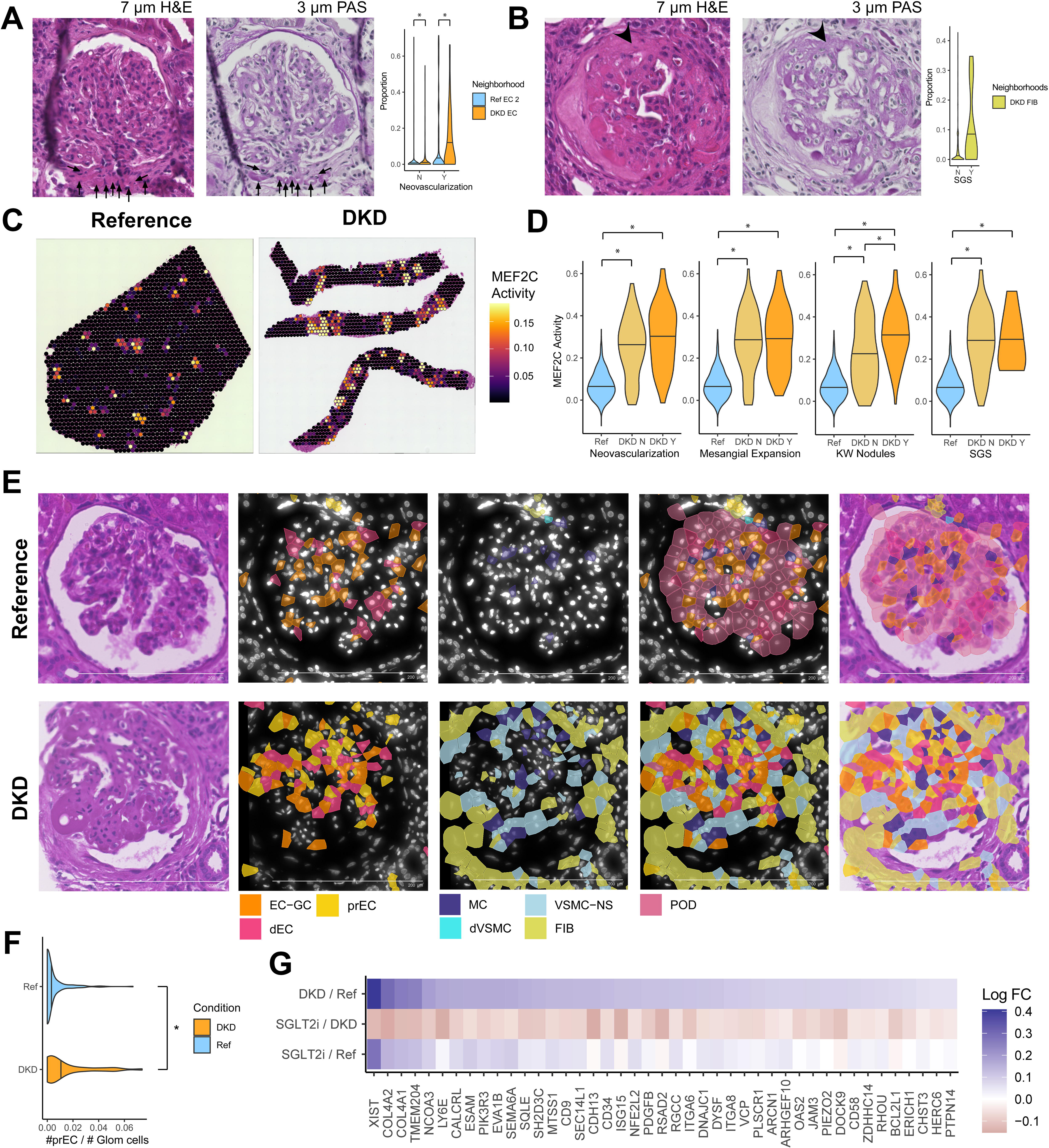
Histopathological and morphological associations with molecular data: **A)** The diabetic kidney disease (DKD) endothelial cell (EC) niche was associated with the presence of histologic neovascularization in glomeruli of FFPE samples. Arrows identify an example of neovascularization near a glomerular hilum. * = p < 0.001. **B)** The DKD fibroblast (FIB) niche was associated with focal segmental glomerulosclerosis (FSGS) lesions in glomeruli of FFPE samples. The arrowhead points to an area of segmental sclerosis as an example. * = p < 0.03. **C)** Transfer score mapping of *MEF2C* activity on FFPE samples reveals localization mainly to glomeruli (but also medullary rays). MEF2C activity was estimated from a vector of composite target gene expression. **D)** *MEF2C* activity was increased in DKD glomeruli and further increased in those with nodular mesangial sclerosis (KW nodules). *= p < 0.01. Ref = reference; DKD N = DKD glomerulus without the histologic feature; DKD Y = DKD glomerulus with the histologic feature. **E)** Example of a reference glomerulus and a DKD glomerulus with neovascularization and nodular mesangial sclerosis with individual cell types annotated. **F)** Ratio of prEC to EC-GC per glomeruli across Ref and DKD samples. * = p < 0.015. **G)** The fold change of selected *MEF2C* targeted genes is altered in DKD and restored after sodium glucose transporter-2 therapy (SGLT2i).

*MEF2C* TF activity was inferred through expression of its target genes in the GRN. Estimated *MEF2C* activity localized over glomeruli and medullary rays in all samples (Figure 6C). However, *MEF2C* had increased activity in DKD glomerular spots. Within DKD glomeruli, *MEF2C* activity was associated with nodular mesangial sclerosis lesions (Figure 6D).

### Spatial localization of EC and VSMC states

With ISS, we identified individual cells and localized the prEC phenotype to DKD glomeruli (Figure 6E). On reference we see the expected glomerular morphology. On a nodular DKD glomerulus, a relative loss of EC-GC was appreciated, with an increase in prEC quantity. In a second glomerulus, MCs transitioned to VSMC-NS, and finally stromal cells along the axis from hilum to nodule (Supplemental Figure S7). A small number of podocytes were seen in both DKD samples. We quantitated the prEC abundance relative to all other cells across each glomerulus in DKD and reference samples. The relative proportion of prECs increased 274.7 ± 1.8% (p = 0.0015) in DKD compared to reference (Figure 6F). Overall, this is consistent with the niches identified in ST. However, the cellular resolution suggests a morphological context to the changes observed in ST and the trajectories identified in multiome.

### SGLT2i therapy reverses the dysregulation of MEF2C activity

To understand the impact of recent therapeutic advances on the *MEF2C* GRN, we accessed a human kidney scRNAseq dataset and assessed *MEF2C* target gene expression in EC-GCs of 3 conditions: healthy kidney, DKD without SGLT2i, and DKD with SGLT2i. In EC-GCs, 226 genes regulated by *MEF2C* were detected. In DKD, 66 genes were differentially expressed at p < 0.05 after Bonferroni correction (64 increased and 2 decreased expression) compared to individuals without DKD. SGLT2i therapy was associated with reversal toward a reference phenotype for 64 of these 66 dysregulated genes, including 43 at adjusted p < 0.05 (Figure 6G).

## Discussion

In this work, we identified a unique niche, with dominant proliferative endothelial cell signature, that was associated with neovascularization in DKD. The niche was characterized by the loss of angiogenic inhibitory communication between the endothelial and VSMC. A TF network composed of *MEF2A, MEF2C,* and *TRPS1* regulated the transition between healthy canonical EC-GCs and proliferative (prEC) or degenerative (dEC) endothelial cell subtypes. *MEF2C* activity and prEC abundance were associated with histologic presence of nodular mesangial sclerosis and neovascularization.

Neovascularization is a complex phenomenon in DKD. It is currently unknown whether neovascularization represents an important adaptation or maladaptation. Glomerular hypertension in DKD may lead to localized hypoxia and sheer stress^50^ and ultimately glomerulomegaly, an early adaptation to DKD, which is followed by subsequent fibrosis^51^.

Hormone and cytokine signaling in DKD may protect from or contribute to neovascularization^52^. VEGF is required to maintain healthy EC-GC and mesangial cell functionality^10, 11^, but the balance of pro-angiogenic and inhibitory processes is tightly regulated. For example, angiopoetin-1 is pro-angiogenic and its interaction with Tie2 (*TEK*) also helps to establish and maintain healthy glomerular vascularity^53–55^. Similarly, *PDGFB*-*PDGFRB* signaling between endothelial and mesangial cells is required for appropriate formation of the capillary network within the glomerular tuft^56, 57^. In contrast, the Notch pathway is well known to inhibit endothelial sprouting through its *DLL4* ligand interaction^42^; however, Jagged1 (*JAG1*) competes with *DLL4* to interact with Notch, restoring a pro-angiogenic state^41^. Consistent with this, in the KPMP dataset, we found maintenance of *ANGPT1-TEK, VEGFA-VEGFRA*, and *PDGFB-PDGFRB* RL pair pro-angiogenic signaling.

*DLL4*-Notch signaling was lost in prECs of DKD subjects, but *JAG1*-Notch signaling was maintained. Traditionally, semaphorins and plexins have been described as contact inhibition proteins in axonogenesis^58, 59^. These proteins have now been shown to be important in the inhibition of neovessel generation^59–61^, but there is limited data in DKD. Our data support the contribution of *SEMA6A* and *PLXNA2* communication between the endothelial and mesangial cell population to maintain a healthy glomerular capillary structure. Our CellOracle KO of the TF network regulating *SEMA6A* and *PLXNA2* predicted a faster progression from an EC-GC to a degenerative endothelial cell and disrupted the transition to a proliferative endothelial cell.

Modelling of single cell trajectories rely on gradual changes in expression between two cell states, and as such, caveats exist. The trajectory may describe an actual transition between a healthy and injured or proliferative cell state.

Conversely, the trajectory may simply represent a spectrum of cell signatures without a direct transition between cell states. Such as distinction is important for the neovascularization in DKD. Possible competing hypotheses include that: 1) proliferation occurs concomitantly from all glomerular endothelial cell subtypes (afferent arterioles, EC-GC, efferent arterioles), or 2) coordinated neovessels emanate from the afferent arteriole of the glomerular hylum. Our data (Supplemental Figure 2 C-F) identified anatomical subclusters within the prEC, which may support either hypothesis. Better mechanistic models might serve to differentiate the origin of prECs.

Our rich dataset employed multiple orthogonal validations. Two disaggregated technologies, snRNAseq and multiome sequencing independently identified a similar set of endothelial subtypes. Likewise, we used three spatial technologies, Visium frozen which held the largest spot sample size, Visium FFPE with improved histologic morphometry, and Xenium which afforded single cell specificity in the histologic context. Combining all these technologies enhanced the confidence of the results. Although we focused our investigation on the glomerular cell cross-talk inherent in neovascularization and nodular mesangial sclerosis, we identified a broad set of molecular changes in DKD. There was substantial overlap in our molecular signature with prior seminal studies. When compared to an atlas of endothelial cell types, our ST glomeruli revealed the expected elevated expression of *PECAM1*, *GJA4*, and *NTN*, but minimal expression of the markers for other ECs^6^. For example, we identified upregulation of genes previously identified in DKD (*IGHM, COL1A2, CXCL6, COL6A3, MMP7*) in the KPMP ST dataset^62^. Further, an analysis of cellular communication in glomerular hyperfiltration identified RL pairs such as *TGFB1*, *VEGFA*, *TGFBR2*, and *PDGFRB*^63^, which aligned with signaling in the KPMP dataset. Thus, our study holds both internal and external validity. In addition, in a study of youth with Type 2 diabetes whose structural changes were limited to minor glomerulomegaly^26^, SGLT2i therapy was associated with a robust reversal of the *MEF2C* activation signatures in glomerular endothelial cells, arguing for the plasticity and reversibility of the pathways described.

While glomeruli adapt to DKD through pro-angiogenic processes, the opposite phenomenon may be observed in the peritubular capillaries of the interstitium. Peritubular capillary rarefaction is a manifestation of chronic kidney disease^64^. The endothelial-to-mesenchymal transition underlying this rarefaction has substantial, but incomplete overlap with the signaling pathways observed in DKD glomeruli. For example, loss of Tie2 and VEGFA signaling accompany this mesenchymal transition. In murine models, conditional deletion of Twist (*Twist1*) and Snail (*Snai1*) both mitigated the endothelial-to-mesenchymal transition in peritubular capillaries. Soluble guanylate cyclase has been proposed as a potential therapeutic option to reduce endothelial dysfunction and prevent downstream proximal tubular epithelial-to-mesenchymal transition^65^.

Our study interrogates the largest ST DKD sample size to date. Despite this, sample size remains a limitation as we were unable to subgroup subjects with DKD according to clinical features. To counterbalance this limitation, we conducted neighborhood analyses on a per spot level and histologic analyses on a per glomerular level to provide our study adequate power. A second limitation is the Visium ST spot size which overlies multiple cells. To overcome this, we included orthogonal validation with ISS, which allows for spatial resolution at the single cell level. Another significant strength of our study was the inclusion of histologic annotations for 264 glomeruli with an adjacent thin section. This strategy enabled us to link the molecular findings to histological features such as nodular mesangial sclerosis and hilar neovascularization.

In summary, we identified a novel proliferative endothelial cell subtype associated with neovascularization in DKD. A common challenge in snRNA-seq analysis is the insufficient power to determine whether two cell subtypes hold distinct biological relevance. Our strategy was to use spatially-defined niches to identify a biologically relevant proliferative endothelial cell cluster. This interrogation method may be applied to other tissues and conditions wherein cellular niches contribute to the molecular characteristics of a cell. Finally, we were able to identify changes in cell-cell communication and the regulatory network underlying the glomerular endothelial cell response to stress in DKD. Our data support that *MEF2C*, *MEF2A*, and *TRPS1* expression and regulation of *SEMA6A* and *PLXNA2* preserve the balance between endothelial cell proliferation and degeneration. In a human scRNAseq dataset, SGLT2i therapy restored the balance of *MEF2C* activity in EC-GCs of DKD towards the reference state in early stage disease suggesting that this pathway can be therapeutically targeted. Studies of pancreas transplantation in type 1 diabetic patients with biopsy proven DKD have shown reversibility of KW nodules^66^. Further investigation is required to understand whether neovascularization may contribute toward resolution or prevention of nodular lesions, or whether therapies should be directed at limiting the extent of angiogenesis.

## Abbreviations

KPMP; HuBMAP; snRNAseq; ST; ISS; snRNA-seq; sc/snRNA-seq; OCT; FFPE; H&E; PAS;

## ACKNOWLEDGMENTS

RMF was supported by a Paul Teschan Research fund Project NO 2023-01 from Dialysis Clinic, Inc. MTE was supported by R01AT011463, R01DK114485, U54 DK134301-01, and U01DK114923. PS was supported by R01DK114485. We would like to acknowledge the NIH common fund Human Biomolecular Atlas Project (HuBMAP) and the NIDDK’s Kidney Precision Medicine Project. This research was supported in part by the Indiana University Pervasive Technology Institute and the Washington University Kidney Translational Research Center. The KPMP is funded by the following grants from the NIDDK: U01DK133081, U01DK133091, U01DK133092, U01DK133093, U01DK133095, U01DK133097, U01DK114866, U01DK114908, U01DK133090, U01DK133113, U01DK133766, U01DK133768, U01DK114907, U01DK114920, U01DK114923, U01DK114933, U24DK114886.

## AUTHOR CONTRIBUTIONS

MTE, RMF, CEA: Conceptualization; SJ, KJK, SER, JRS, RDT: Specimen acquisition; MTE, SJ, RMF, YHC, DLG, CLP, BBL, WSB, FF, MA, AS, DB, MJF, JAS, VN, MK: Data generation; YHC, RMF, DLG, DB: Data curation; RMF, DLG, VN, MTE: Formal analysis; MTE, SJ, PCD, TME, JH, MK: Funding acquisition; All authors: Investigation; All authors: Methodology; RMF: Project administration; RMF, MTE: Writing - original draft; All authors: Writing - review & editing. KPMP: provision of infrastructure.

Please address data and materials requests for Xenium or Visium data to Michael Eadon,

## DATA AVAILABILITY

snRNAseq data was obtained from the KPMP – HuBMAP kidney atlas^14^.Multiome is a published KPMP – HuBMAP dataset^17^. Both datasets, along with the Visium samples, are available on KPMP atlas repository. Code to reproduce analysis can be found on https://github.com/rimelof/Melo_Ferreira_et_al._neovascularization_2024

