## Supplemental Methods for "A *MEF2C* transcription factor network regulates proliferation of glomerular endothelial cells in diabetic kidney disease"

*Visium fresh frozen spatial transcriptomics:* Kidney tissue preserved in Optimal Cutting Temperature compound (OCT) was obtained through the KPMP and processed following a standardized KPMP protocol^1^. Sections of 10 µm thickness were placed on a Visium barcoded slide and stained with hematoxylin (Sigma Aldrich Cat# HHS16) and eosin (Sigma Aldrich Cat# HT110380, H&E). Brightfield images were captured on a Keyence BZ-X810 microscope with a Nikon ×10 CFI Plan Fluor objective. Tissue was permeabilized for 12 minutes and mRNA was captured in the slide’s barcoded spots. The library was prepared following the vendor’s protocol (Visium CG000239 protocol) and sequenced on an Illumina NovaSeq 6000. Reads were aligned to reference genome GRCh38 2020-A and counts matrices were generated with Spaceranger 2.0.0.

*Visium FFPE spatial transcriptomics:* FFPE tissue was obtained from the BBCI^2^. A 7 µm section underwent the Visium FFPE pipeline, while a sequential 3 µm section underwent periodic acid-Schiff (PAS) staining. The 7 µm section was stained with H&E and aligned with a Visium slide using CytAssyst. The tissue was permeabilized and library prepared following the 10x Genomics protocol. The library underwent Illumina sequencing and the resulting reads were mapped to reference genome GRCh38 2020-A with the probeset 2.0 using Spaceranger 2.0.1.

*Single Nucleus RNA sequencing analysis:* The single nucleus RNA sequencing (snRNAseq) dataset was obtained from the published KPMP-HuBMAP atlas^3^. Samples annotated as papilla or medulla were excluded from the analysis. To reflect an inclusive set of cell types potentially localizing to glomeruli, all podocyte (POD), endothelial cell (EC), mesangial (MC) and Vascular Smooth Muscle cell (VSMC) nuclei (N=16,488) were co-clustered and re-annotated from this atlas^3^. Endothelial cells of the glomerular capillary remained annotated as EC-GC. Degenerative endothelial and degenerative peritubular capillary endothelial cells were annotated as dEC, while other endothelial cells (Afferent / Efferent Arteriole, Ascending Vasa Recta, Descending Vasa Recta, Peritubular Capillary, Cycling Endothelial, and Lymphatic) were annotated as Endothelial Cell – Not Specified (EC-NS). Degenerative vascular smooth muscle cells were annotated as dVSMC, while, the remaining vascular smooth muscle cells or pericytes (Renin-positive juxtaglomerular granular cells, vascular smooth muscle cell, vascular smooth muscle cell / pericyte) were annotated as Vascular Smooth Muscle Cells – Not Specified (VSMC-NS). After downstream analyses, a portion of EC-NS was classified as proliferative Endothelial Cells (prEC). A glomerular dataset was generated by subsetting the snRNAseq atlas to include these cell types. Nuclei were clustered correcting for batch effect with Harmony 1.2.0^4^. Cell-cell communications were predicted in the snRNAseq glomerular dataset using the package CellChat 2.1.2^5^

*Visium data analysis:* Within each Visium technology, OCT or FFPE, all samples were merged into a single object. Raw reads were normalized with SCTransform to correct for batch effects^6^. To define glomerular spots, glomeruli were manually annotated using the accompanying histological image and expression of *NPHS2*. Visium spots were assigned to a glomerulus if its center was located inside the annotated glomerulus. Due to the 55 µm diameter of each spot, periglomerular cells were still captured. To remove edge artifacts, the outermost layer of spots on the edge of the samples was excluded. Downstream analyses were conducted on the remaining spots. To estimate cell type composition in Visium Spatial Transcriptomics (ST), the expression of each spot is deconvolved using transfer scores from atlas annotations, calculated using the anchors’ integration method from Seurat 4.3.3^7^.

*Xenium In Situ Sequencing (ISS):* ISS captures RNA transcripts at subcellular resolution, allowing classification of single cells by cell specific markers. FFPE preserved tissue obtained from BBCI or HuBMAP was sectioned and placed on a Xenium slide and processed according to the manufacturer protocol (CG000582 Rev D and CG000584 Rev B). A custom probe set (BMU84Y, Supplemental Table S2) was designed to localize kidney cell types and injury patterns. To obtain histological images, standard staining procedures were followed with H&E staining and destaining, before dehydration and mounting with Eukitt mounting media (Electron Microscopy Sciences Cat #15320). Images were obtained on a Keyence microscope as described above. Histological images were registered to DAPI-stained nuclei images with Xenium Explorer 2.0.0. To identify cells in ISS, a subset of the snRNAseq dataset was mapped with the anchors’ method of Seurat 5.0.3. Epithelial and stromal degenerative cell states were overrepresented due to the limited probe panel and excluded from mapping, along with VSMC-NS. All other glomerular cell type annotations described in the manuscript were maintained.

*Differential Expression and Pathway Enrichment:* Differential gene expression between sets of nuclei or spots was calculated using a Wilcox rank sum test. The significance threshold was set at 0.05 for p values after adjustment with a Bonferroni correction. Pathway enrichment for snRNAseq clusters was performed with SCPA 1.6.1^8^. Single cell groups were defined *a priori* In this pathway analysis method. Then, for each pathway, the cells are clustered according only to the expression of the pathway’s genes, and a q-value is attributed according to the similarity of the clustering to the predetermined groups.

*Cellular niches:* Cellular niches were defined by clustering ST spots according to their cell type composition using traditional Louvain clustering and UMAP projection methods from Seurat. A Fisher’s exact test was utilized to evaluate the likelihood of each cellular niche being overrepresented in DKD samples.

*Annotation of proliferative Endothelial Cells (prEC):* Differential expression between endothelial niches in reference and DKD ST samples identified genes potentially representative of prECs. Reversing the Seurat anchors method which transferred snRNAseq labels to the ST, we next back-mapped the ST niche annotations to the glomerular cells of the multiome dataset to identify an endothelial subgroup preferentially mapping to the DKD endothelial ST niche. This cluster was validated by the expression of differentially expressed genes between the niches. The prEC annotation was then mapped onto the snRNAseq atlas for receptor-ligand modeling.

*Multiome analysis:* The multiome data were obtained from the KPMP-HuBMAP epigenetic atlas^9^. The Seurat object was subsetted to the same cell types as the snRNA-seq dataset with 4410 cells and 983,017 peaks annotated by MACs2. After processing, data were normalized, highly variable features identified, and dimensionality reduction performed through techniques like PCA. The batch effect was removed by harmony (v.1.2.0). The metadata motif database used was JASPAR2022 (v.0.99.7). scMEGA (v.1.0.2) was employed to correlate gene expression with chromatin accessibility at regulatory elements, enabling the modeling of cell trajectories and the identification of regulatory interactions^10^. The *in silico* knockout was modeled with CellOracle (v.0.12.0) using the GRN generated by scMEGA. CellOracle’s simulation tools predicted the downstream effects on target genes and cell states, allowing us to infer the regulatory impact of TF loss^11^.

*Annotation of proliferative Endothelial Cells (prEC):* Differential expression between endothelial niches in reference and DKD ST samples identified genes potentially representative of prECs. Reversing the Seurat anchors method which transferred snRNAseq labels to the ST, we next back-mapped the ST niche annotations to the glomerular cells of the multiome dataset to identify an endothelial subgroup preferentially mapping to the DKD endothelial ST niche. This cluster was validated by the expression of differentially expressed genes between the niches. The prEC annotation was then mapped onto the snRNAseq atlas for receptor-ligand modeling.

*Histopathological scoring of Visium FFPE samples:* Each of the 300 glomeruli in the Visium FFPE dataset was annotated in the sequential H&E (7 µm) and PAS (3 µm) images. Individual glomeruli were scored according to the categorial presence of the following histopathological features: hilar neovascularization; non-nodular mesangial sclerosis (mesangial expansion); nodular mesangial sclerosis (Kimmelstiel-Wilson nodule); and segmental glomerular sclerosis (SGS). To evaluate the association with histopathologic category, a Fisher’s exact test assessed niche mapping proportion and a student’s t-test was used for *MEF2C* activity.

*SGLT2 inhibitor analysis:* The differential expression of genes regulated by *MEF2C* was assessed in the endothelial cluster of a human kidney scRNAseq dataset^12^ for three conditions 1) healthy reference tissue, 2) youth with type 2 diabetes without sodium glucose transporter 2 inhibitor (SGLT2i), or 3) youth with type 2 diabetes with SGLT2i. Fold changes of differentially expressed genes (after multiple-testing adjustment) were depicted in a heatmap.

1.	Michael Eadon RMF, Ying-Hua Cheng, Tarek M. El-Achkar Spatial Transcriptomics Protocol. 2021
