## Supplementary figures and images for "A *MEF2C* transcription factor network regulates proliferation of glomerular endothelial cells in diabetic kidney disease"

### Supplemental Figure 1

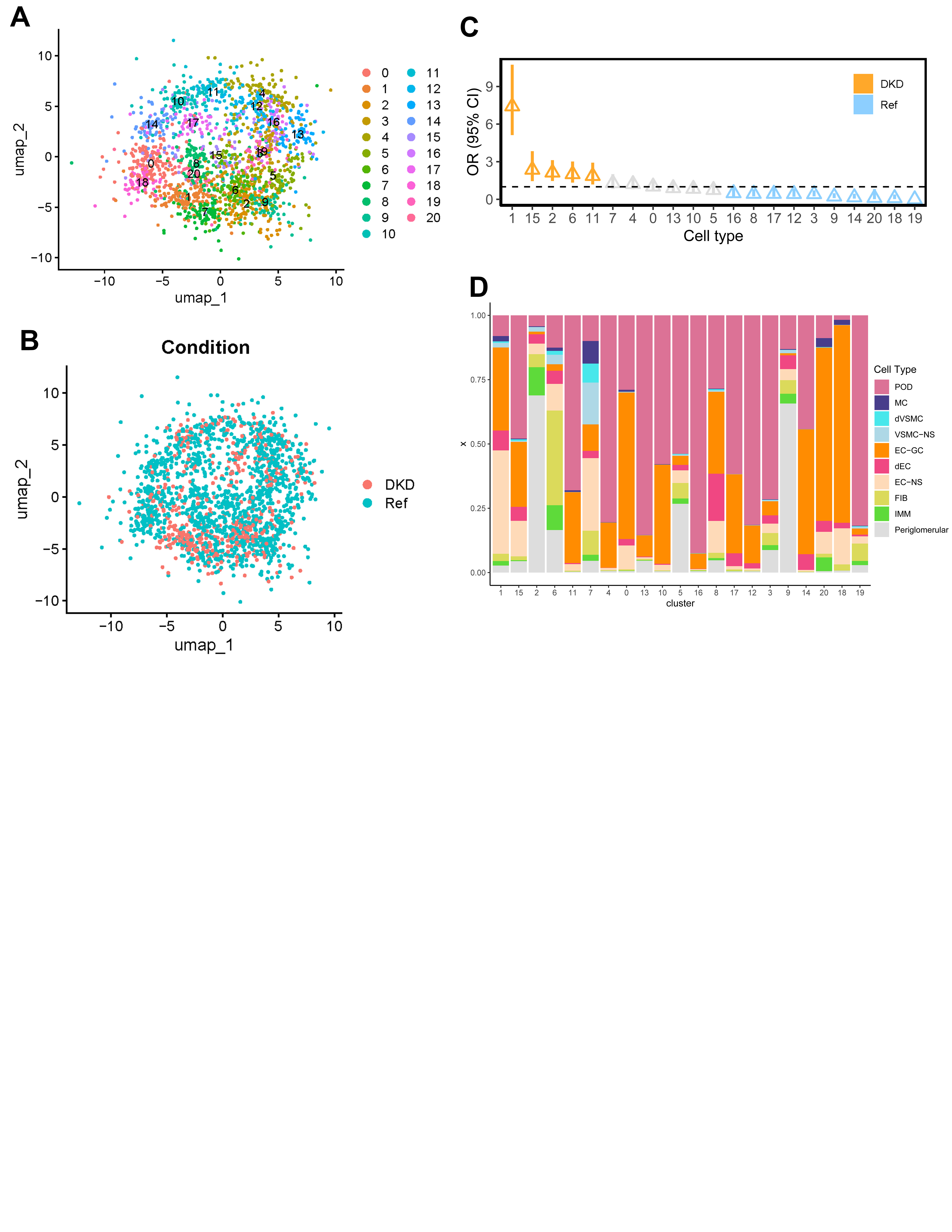

### Supplemental Figure 2

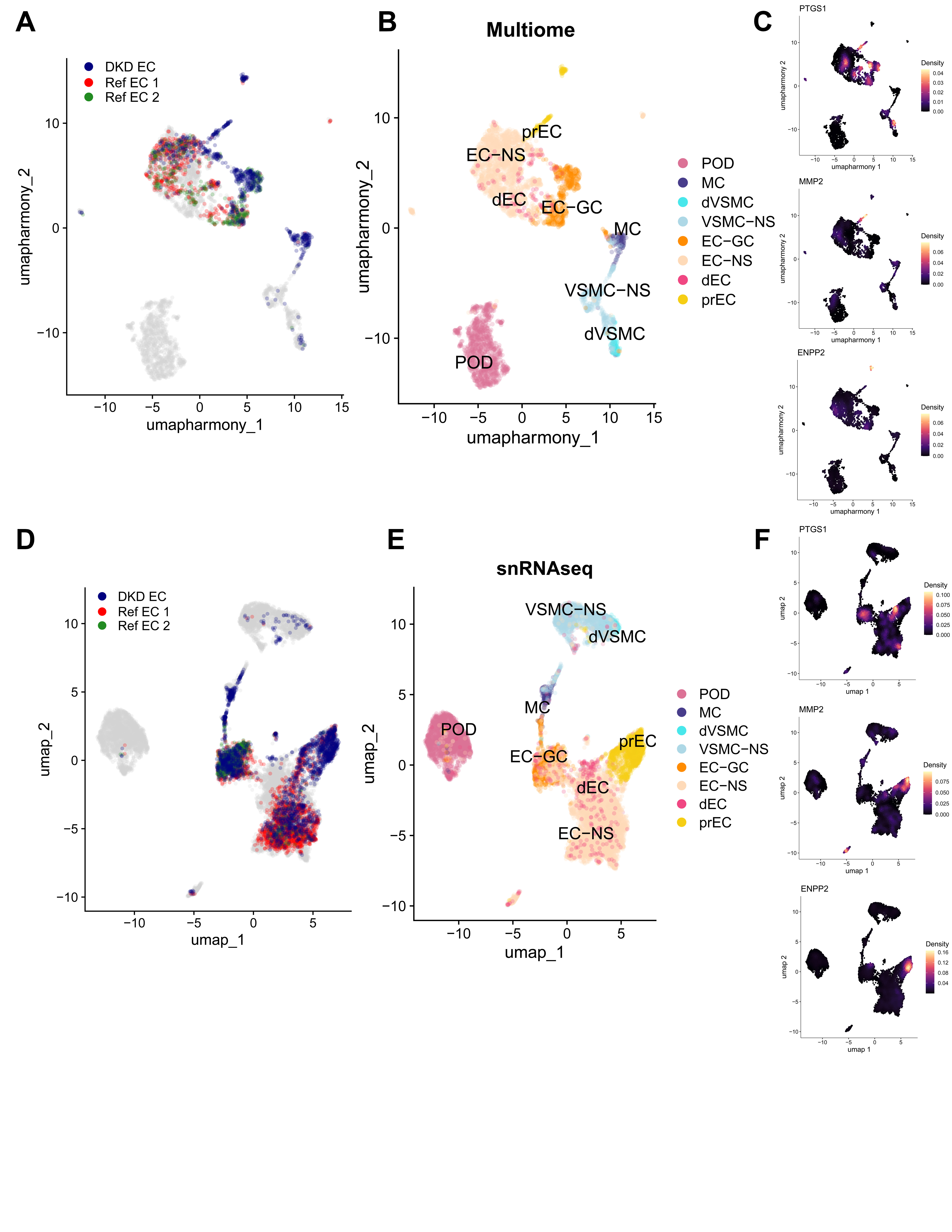

### Supplemental Figure 3

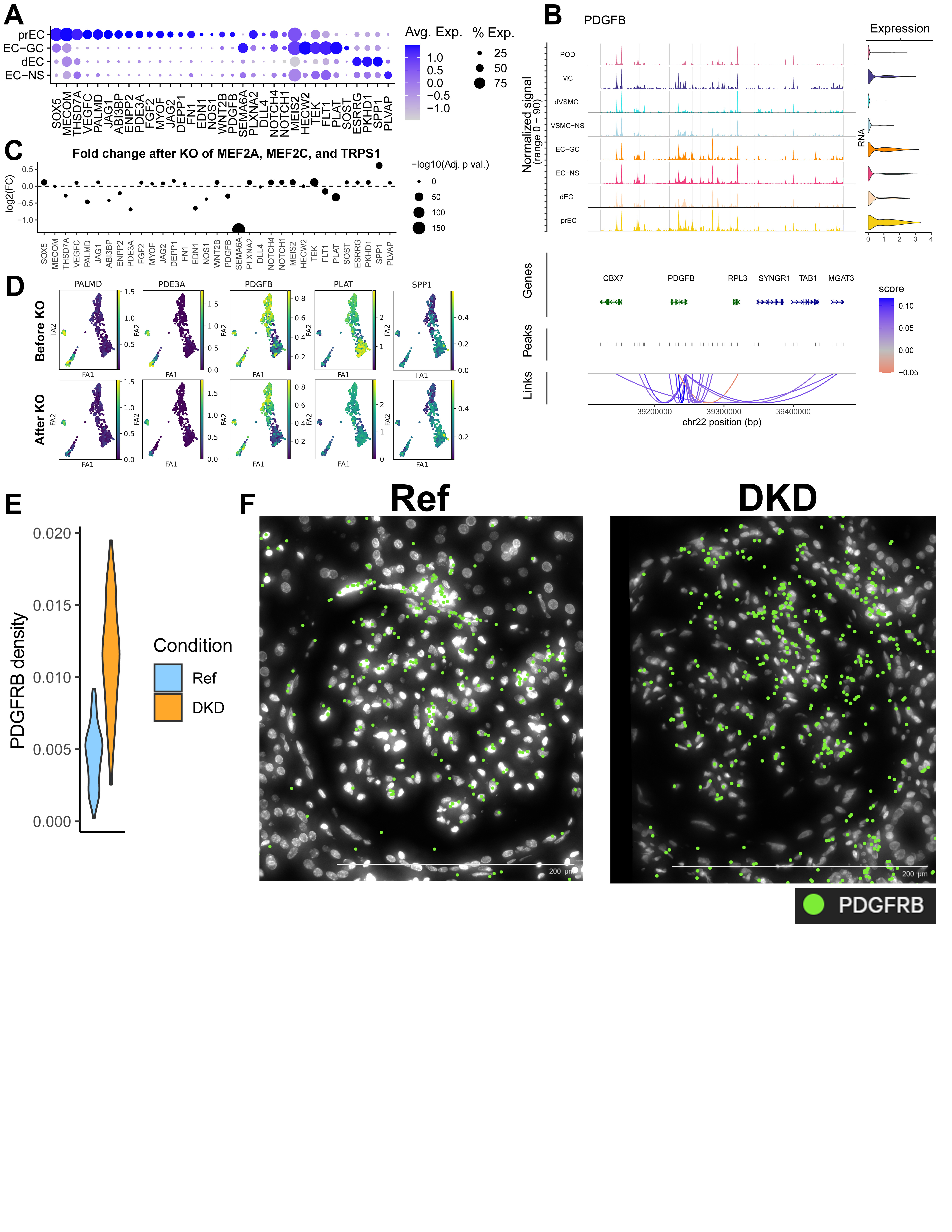

### Supplemental Figure 4

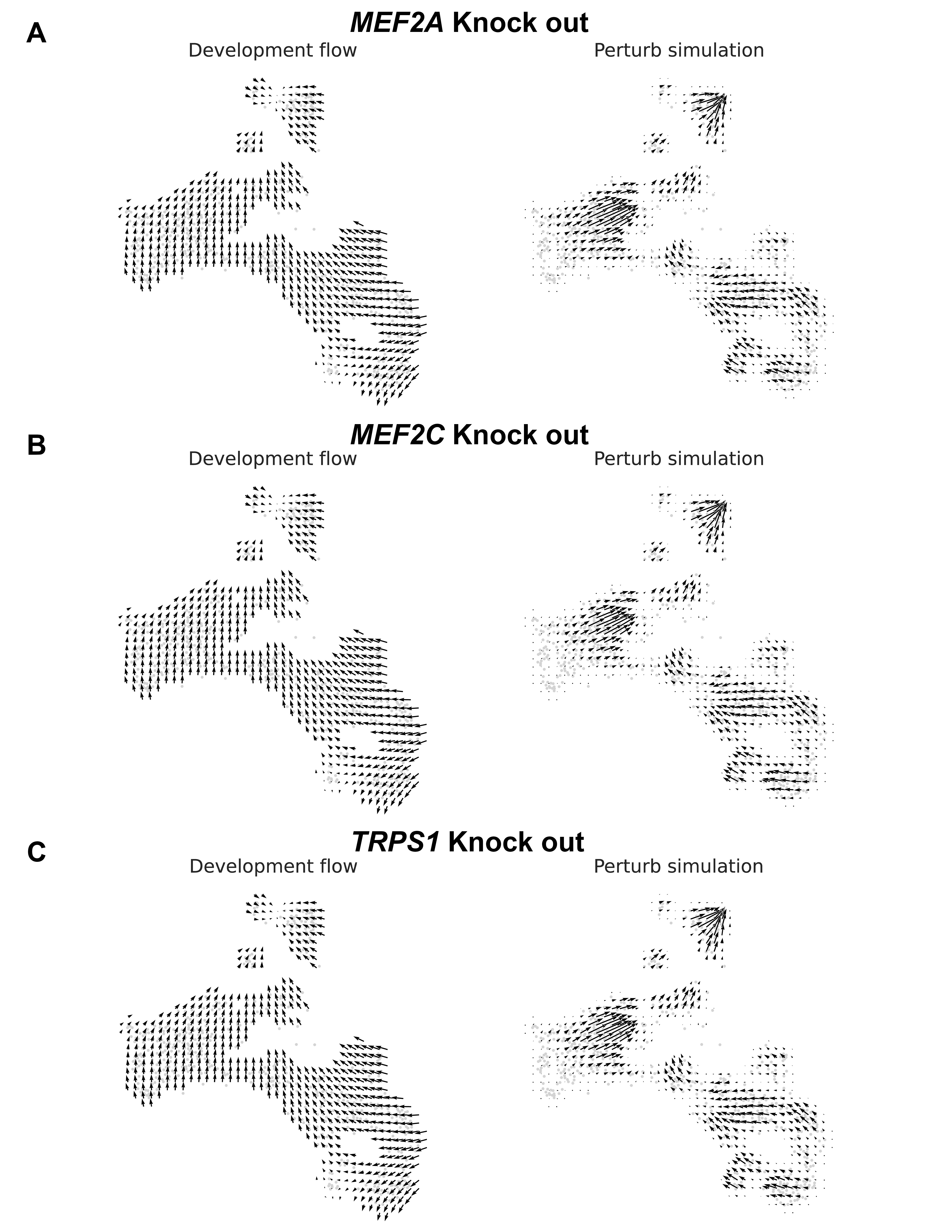

### Supplemental Figure 5

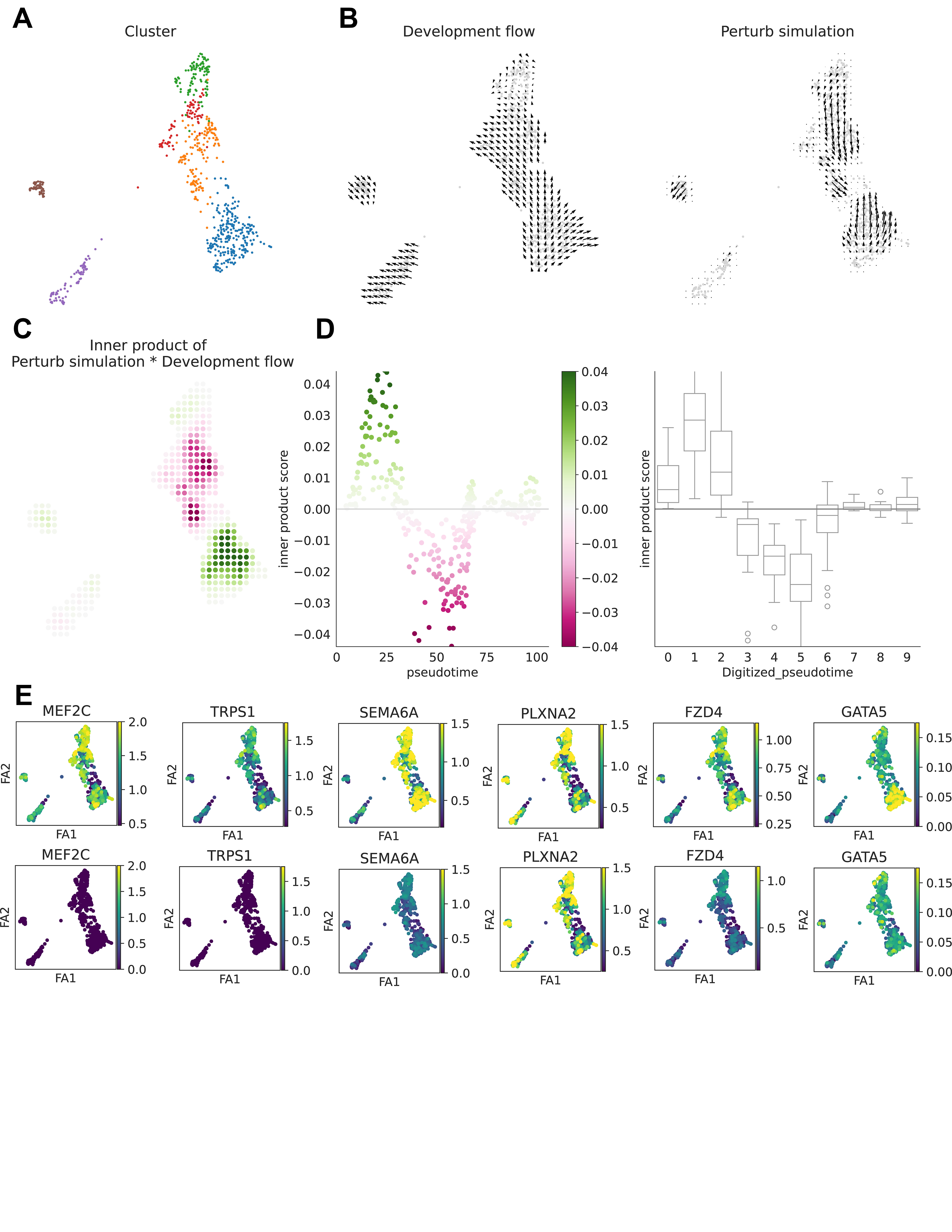

### Supplemental Figure 6

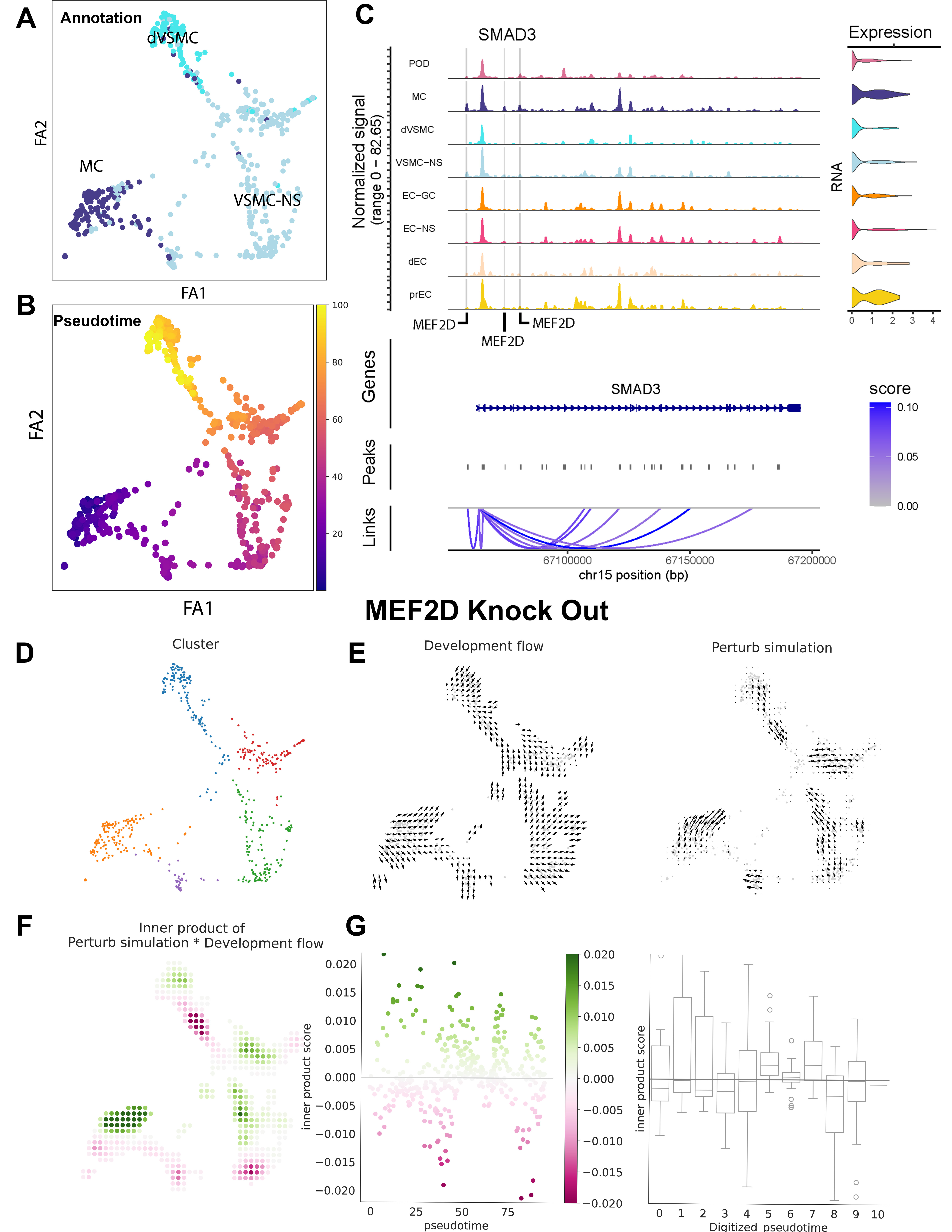

### Supplemental Figure 7

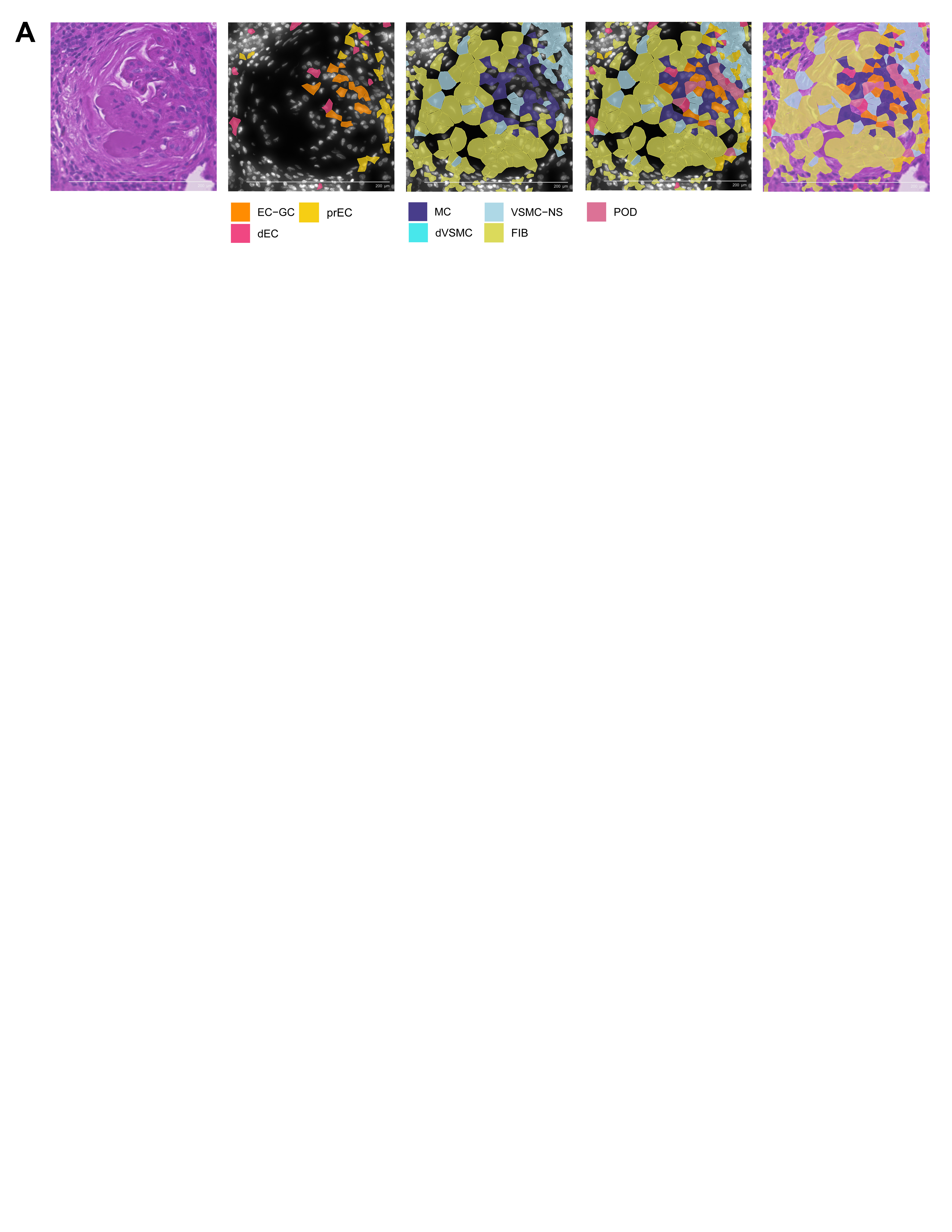
